# Development of lentiviral vectors specifically targeting CD4^+^ cells

**DOI:** 10.64898/2026.11.12.765936

**Authors:** Daniel Cox, Nathalia Almeida, Hataf Khan, Simon Chanas, Michela Marongiu, Conrad Vink, Michael H. Malim, Luis Apolonia

**Author notes:** Correspondence should be addressed to LA.

## Abstract

Because of their central role in immunity, CD4^+^ T cells are a key target for genetic modification in many therapeutic applications, and lentiviral vectors with a VSV-G envelope are often used for this engineering. However, due to their broad tropism, these vectors are inadequate for cell-targeted *in vivo* gene delivery. Here, we explored using HIV-1 Env, which has a natural tropism towards these cells, as an alternative. We describe a three-plasmid vector production system to incorporate a panel of CCR5- or CXCR4-tropic Envs. These Env proteins are expressed from pCAG, a plasmid with reduced HIV-1 elements. We identify HIV-1 Env-incorporating vectors that can be produced with sufficient titres to specifically transduce primary CD4^+^ T cells in the context of *in vitro* mixed populations of peripheral blood mononuclear cells. We show efficient and specific transduction of HIV-1 Env vectors delivering an anti-CD19 chimeric antigen receptor (CAR). We demonstrate functional activity of the generated CD4^+^ CAR T cells, leading to the targeted killing of B cells in those mixed cell populations. Our methodology enables the production of HIV-1 Env-incorporating lentiviral vectors with defined CD4 targeted delivery, opening avenues for direct *in vivo* CAR T cell generation.

## Introduction

Viral vectors are central for the genetic manipulation of cells, with lentiviral vectors being among the most used. These were developed in the late 1990s and utilised in clinical trials from the early 2000s^1,2^. The capacity of these vectors to integrate DNA copies of transduced genes into target cell genomes is especially relevant when prolonged expression is desired in dividing cells, such as haematopoietic stem cells (HSCs) or T cells^3^. With the successful application of chimeric antigen receptor (CAR) bearing T cells to treat cancers clinically, lentiviral vectors have been established as one of the main delivery systems in gene and cell therapy.

CAR T cells are manufactured by isolating T cells from patients, transducing *ex vivo* with an integrating viral vector (based on a g-retrovirus or a lentivirus), expanding and reinfusing them into the patient. Because CAR T manufacturing is a costly and prolonged process, direct *in vivo* transduction would represent an attractive alternative strategy. For this to be successful, the vector would need to specifically target T cells.

One approach to achieve targeted delivery is pseudotyping, where viral cores are coated with the envelope glycoproteins from other viruses, thus altering viral tropism based on the presence of cognate viral entry receptors on target cells. Commonly, lentiviral vectors are pseudotyped with glycoproteins from vesicular stomatitis virus (VSV-G) or from the feline endogenous retrovirus RD114 (RD114), both of which enable the production of high titre vectors under good manufacturing practice conditions^4,5^. These envelopes also broaden the tropism of vector transduction to cells expressing the LDL receptor or a neutral amino acid transporter, respectively, each of which is ubiquitously expressed^6,7^. Critically, this limits their use for *in vivo* cell-selected transduction as targeting is essentially non-specific. Accordingly, there is much interest in developing pseudotyping strategies for targeting specific cell populations, either using natural viral glycoproteins or engineered envelopes containing ligands, single chain antibodies, or antibody mimetic proteins^8^. For example, measles virus and Nipah virus glycoproteins have been engineered to recognise immune cell receptors such as CD3, CD4 and CD8^9–11^.

A viral glycoprotein that naturally targets CD4^+^ cells is the HIV-1 envelope (Env). HIV-1 Env is translated as a polyprotein, gp160, which undergoes glycosylation, oxidation and proteolytic cleavage to yield the transmembrane protein gp41 and the surface-exposed gp120^12–14^. HIV-1 Env is arranged as a trimer on the virion surface^15^. The variable regions of gp120 determine cell tropism, with the V3 loop playing a major role^16–18^ and the V1 and V2 loops also being involved^19^. Env initially binds to CD4, triggering a conformational change that exposes the variable loop regions allowing for co-receptor binding. This triggers membrane fusion, which is driven by gp41^20^. Importantly, co-receptor usage determines which cell populations HIV-1 infects, with CCR5-tropic viruses (R5) broadly infecting macrophages and CD4^+^ T cells and CXCR4-tropic (X4) viruses primarily infecting CD4^+^ T cells^17,18,21,22^. There are also dual-tropic viruses, termed X4R5, which can utilise both co-receptors. Receptor and co-receptor presence and affinity therefore determine which cell subsets are preferentially targeted by HIV-1.

T cell precursors arise from HSCs in the bone marrow, migrating to the thymus where they differentiate and mature^23^. These can be differentiated into CD4^+^ or CD8^+^ T cells, or into non-conventional subsets such as natural T regulatory (Treg) or natural killer (NK) cells. T cell subsets include naïve, memory, and effector cells, which differ in proliferative potential and effector function. Naïve T cells are antigen inexperienced and have low effector function. Memory T cells can be further divided into effector memory (Tem), central memory (Tcm) and stem cell memory (Tscm). Tem cells have low proliferation but high effector function, while Tscm self-renew or differentiate into effector cells, acting rapidly upon antigen stimulation and having high proliferative capacity^23^. For CAR T therapies, preferred targets are less differentiated T cells owing to their proliferation and exhaustion potential, particularly Tscm^24,25^. Importantly, these subpopulations express different levels of CD4, CCR5 and CXCR4^26–28^.

The modular approach to lentiviral vector production (i.e., employing three or four plasmids) enables the routine incorporation of envelope glycoproteins from different viruses. However, to the best of our knowledge, pseudotyping lentiviral particles with HIV-1 Env has only been reported using a two-plasmid system and utilising an Env-expressing plasmid that has additional flanking HIV-1 sequences, thus increasing the potential to generate replication-competent lentivirus through DNA recombination^29,30^. The use of HIV-1 Env in lentiviral vector development has therefore been underexplored. Here, we demonstrate that expression from pCAG, a plasmid that does not contain additional HIV-1 sequences, supports HIV-1 Env expression. This allows for the incorporation of this glycoprotein into HIV-1-based lentiviral vector particles, enabling efficient transduction of cell targets. We further demonstrate CD4^+^ cell-specific delivery and show the functional capacity of transgenes delivered by this vector system, laying the foundation for the development of a targeted CAR T cell therapy.

## Results

### Lentiviral production with a three-plasmid system incorporate HIV-1 Env

Recent advances in CAR T cell technologies have revolutionised the treatment of haematological cancers. Current methodologies typically employ *ex vivo* transduction with CAR-encoding viral vectors pseudotyped with VSV-G. As noted above, the use of VSV-G is unsuitable for *in vivo* transduction strategies because of its broad tropism. In seeking to develop methods for *in vivo* CAR delivery, we therefore explored the use of HIV-1 Env, which utilises CD4 as its receptor. Previous research expressed Env using pSVIII in a two-plasmid system^30^. However, due to the HIV-1 sequences in that expression plasmid (long terminal repeat sequences, and the *tat* and *rev* genes), there is a potential for replication-competent HIV-1 to arise through recombination. Accordingly, we evaluated the production of HIV-1 Env-bearing lentiviral particles using transfected 293T cells and a three-plasmid system that expresses a packaged GFP-encoding indicator genome, a well-characterised HIV-1 Gag and Pol packaging vector, and Env. We compared the transduction efficiency of a HIV-1 Env-containing lentivirus to VSV-G-pseudotyped vectors in TZM-bl cells, a HeLa-based cell line that constitutively expresses CD4, CCR5 and CXCR4 enabling infection with HIV-1 Envs^31^. **Figure 1A** shows that infectious JR-FL Env-bearing vectors are produced by the three-plasmid system, but infectious titres are up to 100-fold lower when compared to vectors pseudotyped with VSV-G.

**Figure 1:**
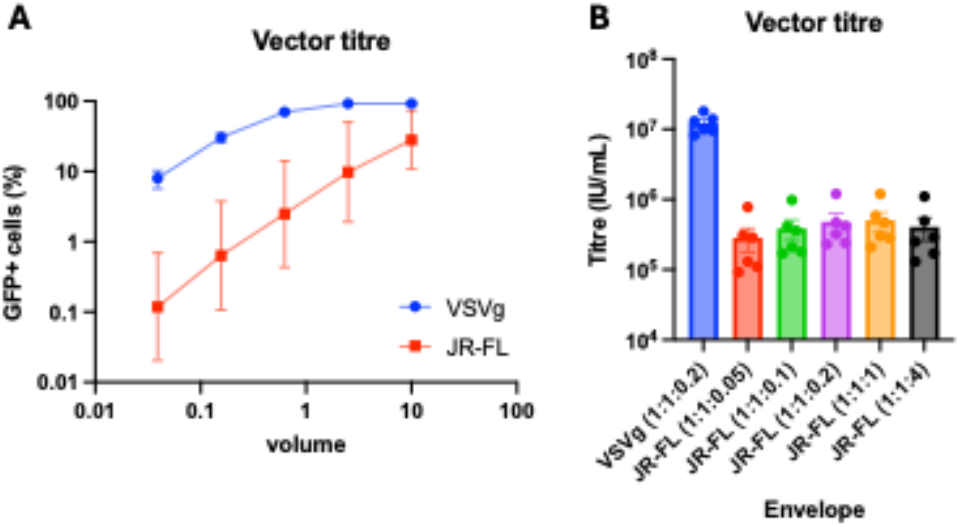
Lentiviral vectors incorporating an HIV-1 Env successfully transduce target ceils. **(A)** Quantification of GFP^+^ TZM-bl cells by flow cytometry. TZM-bl cells were transduced with increasing volume of lentiviral vectors incorporating either VSV-G (blue line) or an HIV-1 Env from the R5-tropic JR-FL isolate (red line) and expressing GFP as a transgene. Graph shows the mean ± SEM. **(B)** GFP reporter transgene expression assessed by flow cytometry. Lentiviral vectors were produced with different ratios of Env-expressing plasmid to packaging and transgene plasmids, and titrated in target cells. Each dot represents the calculated titre for each replicate, with mean ± SEM.

We hypothesised that the lower titres achieved with HIV-1 Env could be due to its inefficient incorporation, so we investigated changing the plasmid ratio during vector production while maintaining the constant amounts of the packaging and genome plasmids. **Figure S1** confirms that the quantity of transfected HIV-1 Env-expressing plasmid correlates with HIV-1 Env protein, while Gag remained unaltered in both producer cells and vectors. **Figure 1B** shows modest vector titres differences, with an optimal vector production in our system been achieved at a ratio of 1:1:1 (packaging plasmid:genome:Env).

### Screen of HIV-1 Env glycoproteins for lentivirus vector production and infection of target cells

We next assessed the functionality of a panel of HIV-1 Env proteins derived from diverse isolates, including R5, X4 and dual-tropic envelopes. Because of possible safety concerns with the pSVIII, we explored alternative expression plasmids. pCAG enabled HIV-1 Env expression in producer cells and incorporation into viral vectors comparable to pSVIII (**Figure S2)**. pCAG expression vector does not harbour any HIV-1 sequence and contains the chicken β-actin promoter with a cytomegalovirus immediate early enhancer element and a chimeric intron (resulting from the fusion between the chicken β-actin intron and a rabbit β-actin splice acceptor) to potentiate high Env expression^32^. Therefore, Env proteins used in this study were expressed from the pCAG plasmid. Activity was measured using stocks generated in transfected 293T cells with the three-plasmid system followed by challenge of TZM-bl cells. **Figure 2A** shows that all Env proteins tested enabled infection of TZM-bl cells, with vectors incorporating the HIV Envs JR-FL, CH040 TF, NL4.3 and 89.6 displaying the highest titres. As noted previously, HIV-1 Envs have tropism for CCR5 or CXCR4 co-receptors or both. We therefore extended beyond TZM-bl cells that express both co-receptors to U87-MG cells ectopically expressing CD4 and either CCR5 or CXCR4. While data obtained in the two U87-MG cell lines yielded similar ranks to TZM-bl cells (**Figure 2B and 2C**), these were not a complete match. Our data illustrate the importance of vector titration in different cell lines when comparing HIV-1 Envs with different co-receptor preferences.

**Figure 2:**
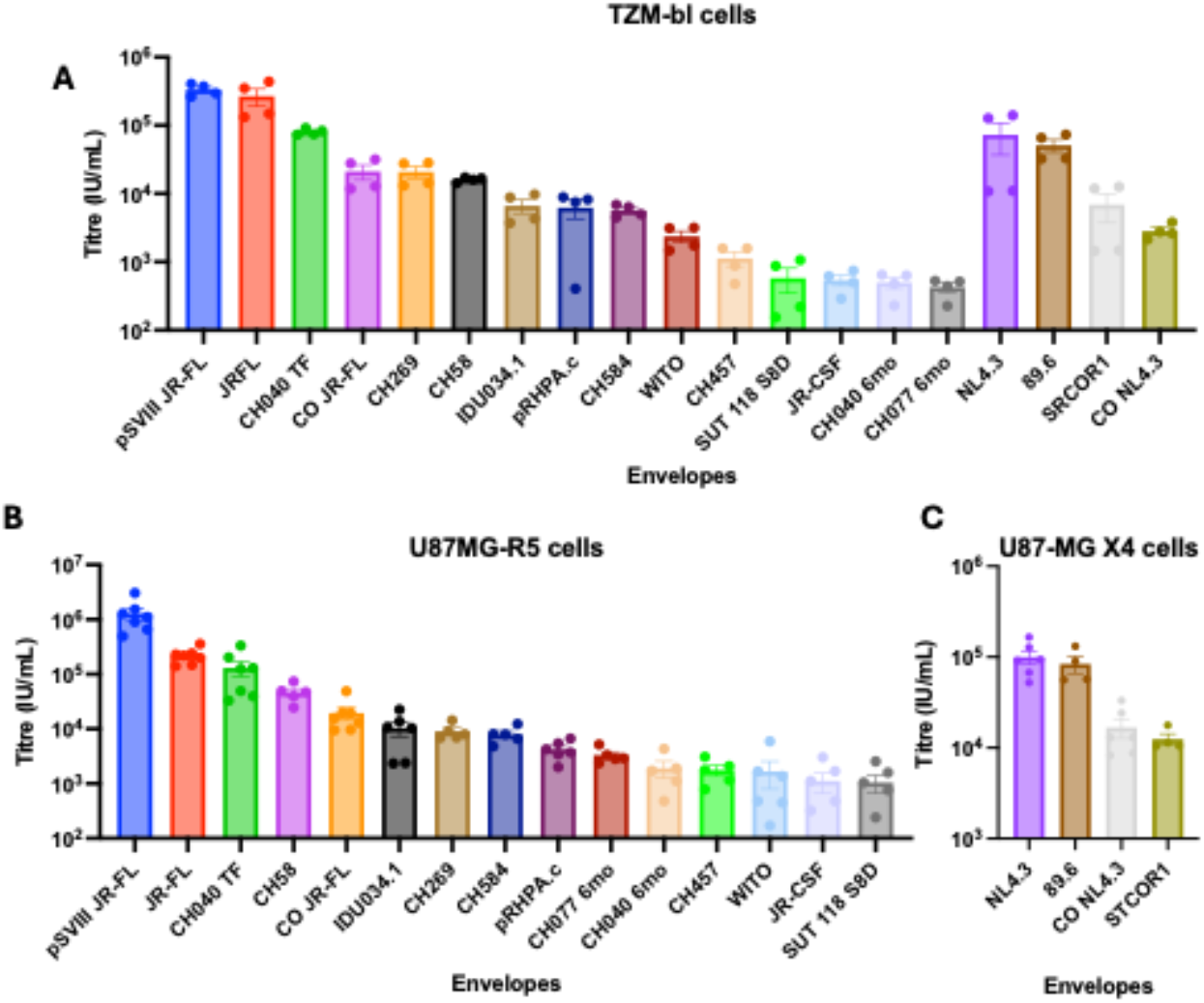
Multiple HIV-1 Env proteins enable vector production and target cell transduction. Titres (IU/mL) of lentiviral vectors produced with a panel of Env proteins from different HIV-I isolates. Titres were calculated from the percentage of GFP^+^ cells measured by flow cytometry 48 hours after transduction of **(A)** TZM-bl cells, **(B)** U87-MG cells expressing CCRS and **(C)** U87-MG cells expressing CXCR4. Graphs show the mean ± SEM. CO: codon optimised.

We show that the R5-targeting Envs JR-FL, CH040TF, and CH58 achieved the highest titres in U87-MG R5 cells. Importantly, the X4 Env NL4.3 and the dual-tropic Env 89.6 were also able to efficiently transduce U87-MG X4 cells.

### Cell specificity of HIV-1 Env-containing lentiviral vectors

Having identified HIV-1 Env proteins that enable the production of vectors, we next investigated their tropism for different lymphocyte populations. We isolated peripheral blood mononuclear cells (PBMCs) from venous blood of healthy donors, activated with CD3/CD28 beads for 24 hours and transduced these cells with a panel of GFP-expressing vectors. Flow cytometry was used to detect specific cell populations and to quantify GFP expression. We also evaluated cell populations pre- and post-activation and confirmed that the distribution of cell types changed within the PBMC population following CD3/CD28 activation: in particular, cells from the myeloid lineage as well as naïve T cells were not detected upon activation^33^, precluding investigation of their transduction (**Figure S4**).

As expected, VSV-G-pseudotyped vectors were able to transduce all lymphocyte populations, including B and NK cells (**Figure 3A**). Importantly, HIV-1 Env-containing vectors were only able to transduce CD4^+^ T cells. Interestingly, in the first three donors tested, NL4.3 Env-pseudotyped vectors achieved an average of 42% infected cells at an MOI of 3 (calculated based on TZM-bl transductions), which was 3 to 6-fold higher than what was achieved with the R5-tropic Envs (**Figure 3A**, closed symbols). On a following experiment (**Figure 3A**, open symbols), we reduced the NL4.3 transducing vectors by 5 fold to an MOI of 0.6 and included 89.6 Env-pseudotyped vectors, also at MOI 0.6. Nonetheless, we obtained a similar transduction level of CD4^+^ T cells compared to the other vectors tested (calculated to be 1.5 for NL4.3 and 0.8 fold for 89.6 vectors, on average). Our results highlight the challenges in comparing functional titres of R5- and X4-tropic vectors side-by-side, likely due to variations in relative expression of CCR5 and CXCR4 proteins across different cells.

**Figure 3:**
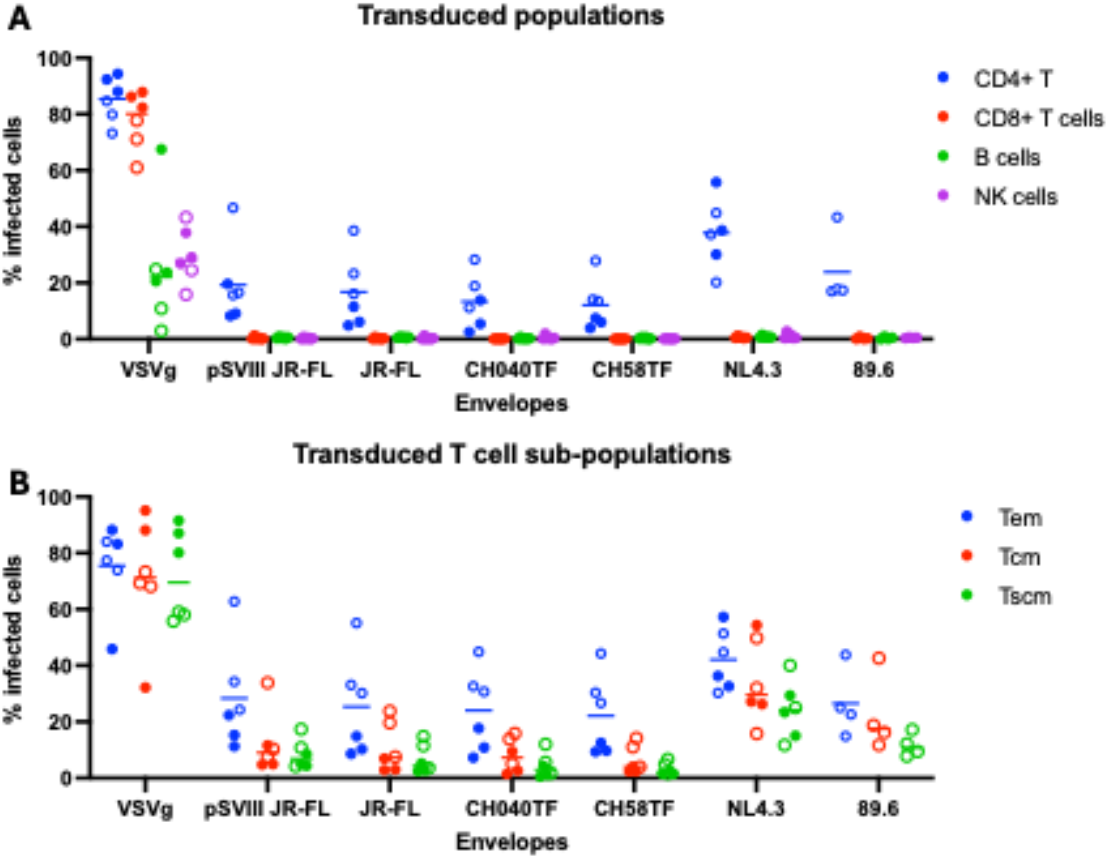
HIV-1 Env-containing vectors transduce CD4^+^ T cells specifically. Percentage of GFP^+^ PBMCs following transduction at MOI 3 with lentiviral vectors expressing a OFP reporter gene and incorporating VSV-G or a panel of HIV-1 Envs. **(A)** Transduction of PBMC populations: CD4^+^ T cells, CDS^+^ T cells, B cells and NK cells. **(B)** Transduction of CD4^+^ T cell subsets: effector memory (Tem), central memory (Tcm) and stem cell memory (Tscm). Each dot represents one donor and bars the mean. PBMCs were activated with CD3/CD28 beads for 24 hours prior to transduction. After 72 hours, cells were stained with a live/dead dye and cell-surface markers, and populations were identified by flow cytometry. Open circles for NL4.3 and 89.6 Env vectors indicate transduction at MOI 0.6.

As outlined above, circulating T cells can be divided into subpopulations, depending on their state of differentiation or activation. Broadly, activated T cells are short-lived, but can also persist as memory T cells, and provide a faster (memory) response if the same antigen is detected. We therefore assessed the susceptibility of CD4^+^ memory T cell subpopulations to transduction by the vector panel used in **Figure 3A**. As seen with bulk CD4^+^ T cells, the vector carrying the X4-tropic NL4.3 Env transduced Tem, Tcm and Tscm cells more effectively than the other Envs (**Figure 3B**). The dual-tropic 89.6 Env enabled transduction of all CD4^+^ memory T cell subpopulations, with Tcm and Tscm infections also being notably more efficient than the R5-tropic Envs. Indeed, for all R5-tropic Envs, transduction of Tem cells was markedly more efficient than for Tcm and Tscm cells, which is in accordance with previous publications, a phenotype postulated to be due to higher levels of CCR5 expression in the Tem cells^26–28^.

### Delivery of CD19-targeting CAR to T cells using HIV-1 Env lentiviral vectors

Having demonstrated that HIV-1 Env-containing vectors can transduce primary CD4^+^ T cells, we next explored whether our approach could deliver a clinically relevant chimeric antigen receptor (CAR) transgene designed to recognise CD19, an established therapeutic target in B-cell malignancies^34^. Based on our results to date, we focused our efforts on the JR-FL and NL4.3 Env proteins. We transduced CD4^+^ T cells isolated from the PBMCs of three donors with vectors encoding either GFP or the CAR. GFP expression was detected as before, and the CAR was detected using a FITC-labelled CD19 protein (this allows identification of cells expressing the correctly folded CAR).

**Figure 4A** demonstrates that all vectors were able to transduce these CD4^+^ T cells, and that the CAR can be detected by flow cytometry. As expected, VSV-G conferred the highest levels of transduction, followed by NL4.3 Env and JR-FL Env. All cell populations were then cultured together with Nalm6 targets (CD19-expressing cells) and CEM-SS non-target (which do not express CD19) cancer cell lines. Target and non-target cancer cells had previously been modified to express luciferase, which enabled direct quantification of surviving cells after co-culture, and therefore, measurement of CAR-mediated cell killing. **Figures 4B and 4C** show that GFP-expressing cells do not kill either target or non-target cells, as luciferase values were similar to cultured target cells alone. In contrast, CAR-expressing CD4^+^ T cells were able to kill Nalm6 target cells but not CEM-SS cells, indicating that the CAR specifically recognised CD19 and triggered targeted cytotoxicity. Similar results were obtained with a different effector:target ratio (**Figure S3**).

**Figure 4:**
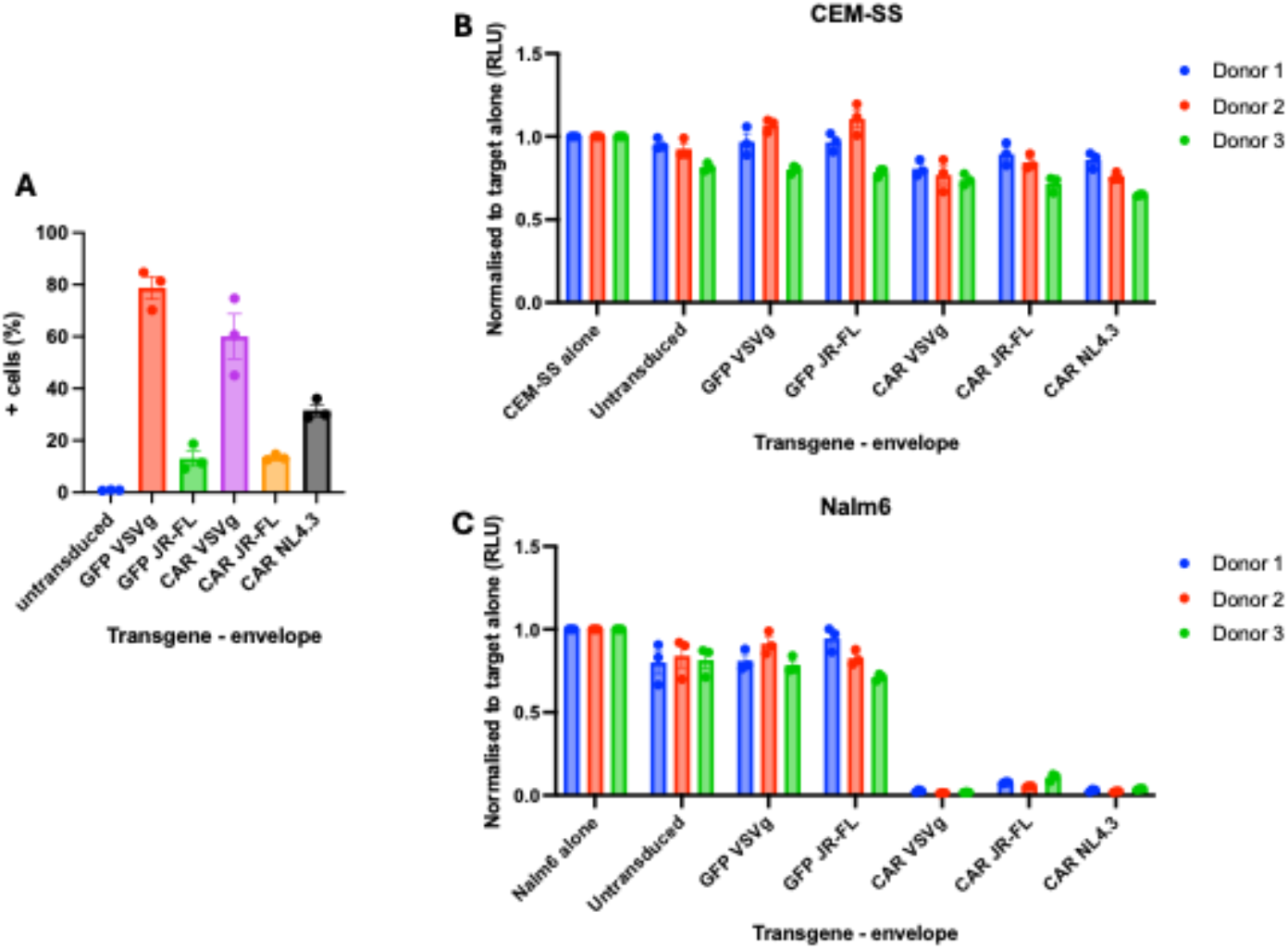
Primary CD4^+^ CAR T cells kill target CD 19^+^ cancer cell lines. **(A)** Transgene expression in primary CD4^+^ T cells transduced with vectors incorporating VSV-G or the indicated HIV-1 Env and expressing either GFP or a CAR targeting CD 19. CD4^+^ T cells were isolated and activated prior to transduction. Percentage of cells expressing GFP and CAR (stained with a CD 19 antibody conjugated with FITC) was measured by flow cytometry. Each dot represents one donor. **(B, C)** Killing of luciferase expressing **(B)** CEM-SS cells (CD19 negative, non-target control) and **(C)** Nalm6 cells (CDI9^+^ target) by CAR-transduced CD4^+^ T cells, cocultured at an effector: target ratio of 1:1 for 48 hours. Surviving cells were measured by luciferase activity and normalised to target cells alone. Graphs show mean ± SEM.

We next hypothesised that our HIV Env-incorporating vectors could deliver the CD19 CAR specifically to CD4^+^ T cells within mixed primary PBMC, resulting in the direct killing of B cells within that population. For this, we transduced whole PBMC cultures with the anti-CD19 CAR-expressing vectors. **Figure 5A** confirms that the VSV-G-pseudotyped vector delivers the CAR transgene to all cell types, whereas only CD4^+^ T cells express the CAR when HIV-1 Env-carrying vectors are used. Transduced PBMCs were then cultured for 72 h and distinct cell populations quantified by flow cytometry. In addition to the CAR, we also used the GFP-expressing vector as a control. B cells could be cultured in the presence of GFP-expressing cells. In contrast, we observed a dramatic reduction in the number of B cells in the presence of CAR-expressing cells (0.02% versus 1.5% of total cells, respectively), indicating that B cell-cytotoxicity was CAR-dependent (**Figure 5D)**. In addition, we observed that other cell populations, such as CD4^+^ T, CD8^+^ T or NK cells did not change in frequency when compared with GFP-transduced PBMCs, and only the B cell population was depleted, demonstrating the selectivity of the CAR towards CD19-expressing cells (**Figure 5B-5E**). Taking these results together, we conclude that B cells within PBMC cultures are specifically depleted following transduction of the whole PBMC population with vectors expressing a CD19-targeting CAR, and that HIV-1 Env-carrying lentiviral vectors can mediate this action through the specific transduction of CD4^+^ T cells.

**Figure 5:**
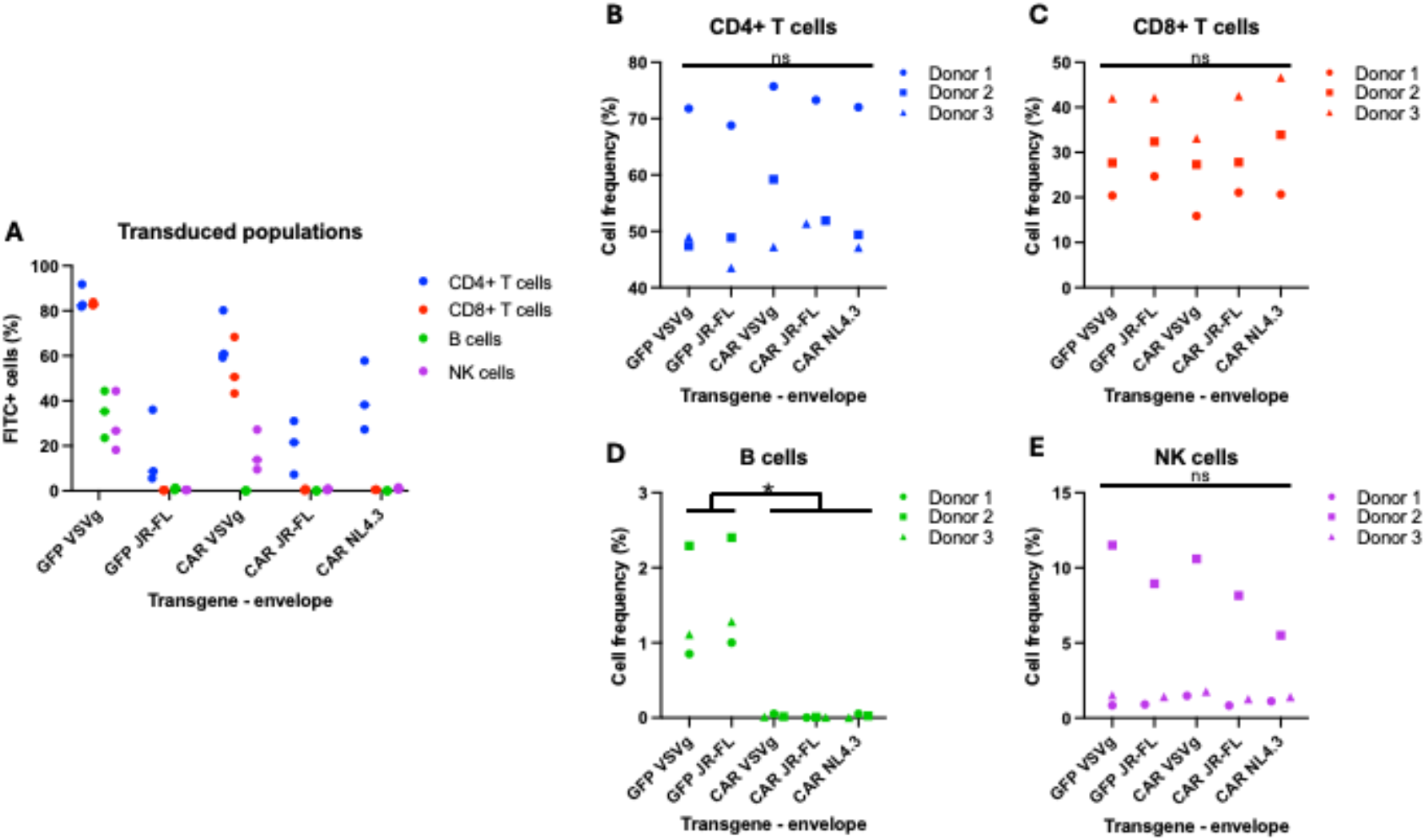
CD4^+^ CAR T cells generated ill whole PBMCs specifically deplete the B cell population. **(A)** Percentage of transgene expressing cells in PBMC mixed population following transduction with vectors pseudotyped with VSV-G or the indicated HIV-1 Env, and expressing either GFP or an anti-CD19 CAR. GFP was detected directly and the CAR with a FITC-conjugated CD19 protein using flow cytometry. Each dot represents one donor**. (B-E)** Frequency of **(B)** CD4^+^ T cells, (**C)** CDS^+^ T cells, **(D)** B cells and **(E)** NK cells, as a percentage of total PBMCs. Each symbol represents one donor. PBMCs were activated with CD3/CD28 beads for 24 hours, transduced, and analysed 72 hours later using antibodies to identify specific populations by flow cytometry. Statistical significance was determined by one-way ANOVA followed by Tukey’s multiple comparisons test; * *p <* 0.05; ns, not significant.

## Discussion

Lentiviral vectors are routinely pseudotyped with a glycoprotein envelope from a different virus, such as VSV-G. This glycoprotein confers a broad vector tropism because the receptor for this viral envelope protein, the LDL receptor, is ubiquitously expressed. In addition, the stability of this envelope facilitates vector production and storage at scale^4,7^. However, this technology is not well suited to applications in which specific cell lineages are to be targeted by vectors. Due to their key role in immunity, T cells are of particular interest for genetic manipulation. A clinically relevant example is T cell genetic alteration to achieve sustained CAR expression. CARs enable the transduced cells to recognise specific proteins on target cells, such as cancer cells, triggering a signalling cascade that leads to the elimination of tumour cells^35^. The clinical success of CAR T cell therapy has resulted in seven CAR T cell therapies approved by the Food and Drug Administration since 2017, with additional CAR T cell products and next-generation platforms currently under clinical development^36,37^.

Current CAR therapies involve the use of retroviral vectors to transduce T cells *ex vivo*^36,38^. Nonetheless, using vectors to generate CAR T cells *in vivo* would considerably reduce the manufacturing complexity, enable an off-the-shelf viral vector product, with shorter turnaround times and simpler manufacturing, with a marked lower cost for these therapies^39^. However, vectors harbouring envelopes with broad tropism, such as VSV-G, are not conducive for cell-targeted *in vivo* systemic delivery, and technologies that target specific cells need to be developed^10,39,40^.

To target vectors towards T cells, we used the envelope of HIV-1, which is known to depend on CD4 and a co-receptor, such as CXCR4 or CCR5 for entry into cells^15,17,18,22^. We utilise a three-plasmid system that minimises the production of replication-competent recombinants^1,2,41^. Using the well-characterised Env of the JR-FL isolate, we show that, despite obtaining a lower titre compared to VSV-G-pseudotyped vectors, HIV-1-based vectors incorporating this HIV-1-derived Env efficiently transduced cells that express CD4 and CCR5. Moreover, optimising the ratio of envelope plasmid during transfection increased vector production. Env modifications that prevent gp120 shedding or stabilise this protein^42^ in vector particles could be investigated to further improve vector titres.

We then evaluated a library of HIV-1 Envs from different viral isolates, expanding the number of HIV-1 Envs that can be incorporated into lentiviral vectors. These included HIV-1 Envs that preferentially target CXCR4, CCR5 or either co-receptor. When vectors with these Envs were tested in PBMCs, we confirmed the specific tropism for T cells. Nonetheless, due to the activation protocol used, we were unable to evaluate the tropism for cells from the myeloid lineage^33^.

It is known that CAR T cells can become exhausted and may not persist for prolonged periods^43,44^. Central memory (Tcm) and stem cell memory (Tscm) T cells are known to have a high proliferative capacity while maintaining effector functions, and these have been shown to be preferable starting populations to generate CAR T cells^24,25^. Here, we observed that X4-tropic Envs, such as NL4.3, transduced Tscm and Tcm cells with higher efficiencies than R5-tropic Envs, while maintaining a high transduction efficiency in T effector cells. This observation is consistent with the differential HIV-1 co-receptor expression across T cell subpopulations shown previously, including CXCR4 expression in less differentiated T cell populations and increasing CCR5 expression following T cell differentiation^26–28^. Taken together, our results highlight X4-Envs as a promising avenue for future development aimed at targeting Tscm cells for CAR expression.

To evaluate the delivery of a clinically relevant transgene, we used a CAR that recognises the therapeutic target CD19^34^, expressed in B cells. PBMCs transduced with vectors expressing this CAR were able to kill cancer cells that express the CD19 protein. In addition, we demonstrated the efficacy of HIV-1 Env-incorporating vectors in the context of mixed populations. T cells were specifically transduced by the vectors, leading to the complete elimination of B cells, the population of cells in PBMCs that express the CD19 CAR target.

Overall, we demonstrate that HIV-1 Env-pseudotyped lentiviral vectors specifically transduce CD4^+^ T cells and mediate expression of a therapeutically relevant CD19-targeting CAR, resulting in selective elimination of B cells in primary PBMC cultures. Together, these findings lay the foundation for the investigation of HIV-1 Env-containing lentiviral vectors in *in vivo* models and open avenues for future gene therapies.

## Material and Methods

### Plasmids

Plasmids used in lentiviral vector production were described before and included pCMV-ΔR8.91 (p8.91), pHRsin.SFFV.eGFP, pMD2.G (that expresses VSV-G)^2,41^, and pSVIII JR-FL^45^. HIV-1 gp160 Env cDNAs were PCR-amplified with primers containing the recognition sequences of the restriction enzymes EcoRI and NotI and ligated to the pCAG plasmid^32^, modified to contain EcoRI and NotI between the promoter and polyA signal. Codon-optimised (CO) JR-FL and NL4.3 Env cDNAs were used where indicated. We used a second-generation CAR comprising a CD8α leader sequence, the FMC63 anti CD19 scFv, a CD8α hinge and transmembrane, a 4-1BB co-stimulatory domain and the CD3ζ signalling domain. This protein was expressed from a EF1α promoter.

### Cells

Human embryonic kidney (HEK) 293T (ATCC), Uppsala 87 Malignant Glioma (U87-MG, ATCC) CXCR4^46^, U87-MG CCR5^46^ and TZM-bl cells (NIH AIDS Reagents Repository Program) were maintained in Dulbecco’s modified Eagle’s medium (DMEM) with GlutaMAX (Thermo Fisher Scientific) supplemented with 10% heat-inactivated foetal bovine serum (FBS) and 1% penicillin-streptomycin (P/S) (Invitrogen). Raji, Nalm6 (ATCC) and CEM-SS cells (NIH AIDS Reagents Repository Program), stably transduced with a lentiviral vector expressing luciferase, were cultured in Rosewell Park Memorial Institute (RPMI) with GlutaMAX and 25 mM HEPES (Thermo Fisher) supplemented with 10% heat-inactivated FBS and 1% P/S. Cell lines were routinely screened for mycoplasma contamination using the MycoAlert Mycoplasma Detection Kit (Lonza).

PBMCs were isolated from venous whole blood collected from consented healthy donors through the King’s College London Infectious Diseases BioBank (ethics reference LA-241121) under overall permission from the Southampton and South West Hampshire Research Ethics Committee (B) (REC reference 19/SC/0232). PBMCs were separated by density gradient centrifugation using SepMate and Lymphoprep™ (STEMCELL Technologies), according to the manufacturer’s protocol. CD4^+^ T cells were isolated from the PBMC population using negative cell sorting with the CD4^+^ T cell isolation kit, human (Miltenyi Biotec), following the manufacturer’s instructions. Cells were grown in RPMI with GlutaMAX and 25 mM HEPES, supplemented with 10% heat-inactivated autologous serum, 1% P/S and 30 U/mL rIL2 (Roche). Prior to transductions, cells were activated with dynabeads human T-activator CD3/CD28 beads (Thermo Fisher) at a 1:1 cell:bead ratio.

### Vector production and titration

Lentiviral vectors were produced by co-transfecting HEK293T cells with plasmids p8.91, the transgene plasmid and the Env plasmid at a ratio of 1:1:1 (unless otherwise stated) using transIT-2020 (Mirus). Supernatants were collected 48 hours post-transfection, filtered through a 0.45 µm membrane, and where stated, concentrated by centrifugation through a 20% weight/volume (w/v) sucrose cushion at 4800 x *g* for 18 hours at 4 °C.

GFP-expressing vectors were titrated in TZM-bl cells and where stated, also in U87-MG cells. Specific volumes of vectors were added to 1x10^4^ target cells and incubated for 48 hours. Cells were then harvested and GFP+ cells were assessed by flow cytometry on a BD FACSCANTO II flow cytometer. Titres were calculated based on TU/mL = (% GFP^+^ cells/100) × (number of infected cells) / volume of vector stock added (mL).

### Flow cytometry

Adherent cells were detached with TrypLE (Gibco) or 6 mg/mL lidocaine (Sigma-Aldrich, L5647) and 10 mM EDTA (Gibco) in PBS. PBMCs were incubated with Zombie™ NIR live/dead dye (Biolegend) and stained with conjugated antibodies: CD3-BUV395 (UCHT1, Becton Dickinson), CD4-Alexa Fluor 700 (SK3, BioLegend), CD8a-PerCP-Cy5.5 (RPA-T8, BioLegend), CD16-BUV737 (3G8, Insight Biotechnology), CD56-PE-Texas Red (HCD56, BioLegend), CD19-Brilliant Violet 711 (HIB19, BioLegend), CD14-Brilliant Violet 605 (M5E2, BioLegend), CD197-Brilliant Violet 421 (G043H7, BioLegend), CD95-PE-Cy7 (DX2, BioLegend), CD45RA-Brilliant Violet 510 (HI100, BioLegend), CD45RO-APC (UCHL1, BioLegend), CD25-PE (BC96, BioLegend), CD127-Brilliant Violet 785 (A019D5, BioLegend). CAR molecules were detected with FITC-labelled Human CD19 (20-291, (ACROBiosystems). All samples were analysed on a CytoFLEX Flow Cytometer (Agilent), and data were processed using FlowJo software (version 10).

### Cell line cytotoxicity assay

Target or non-target cell lines expressing luciferase were co-cultured for 72 hours with or without CAR-expressing CD4^+^ T cells at 1:1 or 1:10 effector:target ratio. Luciferase expressed by surviving cells was measured with OneGlo assay (Promega) following manufacturer’s instructions, with luminescence assessed on a PerkinElmer Victor X Light luminometer.

### Statistical tests

Statistical analysis was performed with GraphPad Prism (version 10). Individual data points, mean and SEM are shown in graphical representations. The statistical test employed in Figure 5 was a one-way ANOVA, followed by Tukey’s multiple comparisons test. Statistical significance was set at p<0.05 (*).

## Data availability

The data and materials that support the findings of this study are available from the corresponding author upon reasonable request.

## Acknowledgements

The authors would like to thank Dr. Anna Rose, Advanced Cytometry Platform at Guy’s and St. Thomas’ NHS Foundation Trust, for support and helpful discussions and Stelios Papaioannou for the provision of reagents. This work was funded by a BBSRC industrial Collaborative Award in Science & Engineering (iCASE) grant in partnership with GlaxoSmithKline (BB/V509632/1). N.A. was funded by the Wellcome Trust PhD programme in Cell Therapies and Regenerative Medicine (108874/Z/15/Z). M.H.M is funded by the Wellcome Trust (222433/Z/21/Z).

## Author contributions

C.V. and L.A. conceived the study. D.C., M.H.M., and L.A. designed the experiments. S.C., M.M., and C.V. contributed to experimental design. D.C., N.A. and HK performed the experiments, collected and analysed data. M.H.M. and L.A. directed the study. L.A. wrote the manuscript with input from all authors.

