## Supplemental Data for "Development of lentiviral vectors specifically targeting CD4^+^ cells"

### Supplemental material

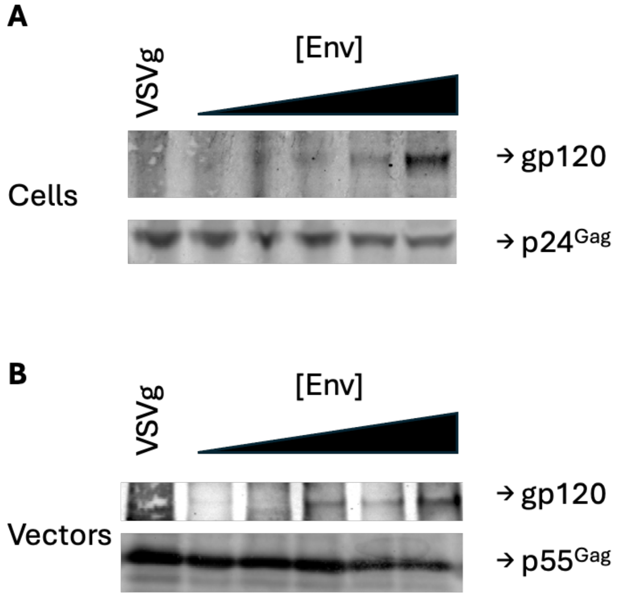


**Figure S1: HIV-1 Env expression and incorporation correlates with quantity of plasmid transfected.**
HEK 293T cells were transfected with varying doses of Env-expressing plasmid (as described in Figure 1. HIV-1 Env and Gag were analysed by immunoblot in **(A)** producer cells and **(B)** vectors.

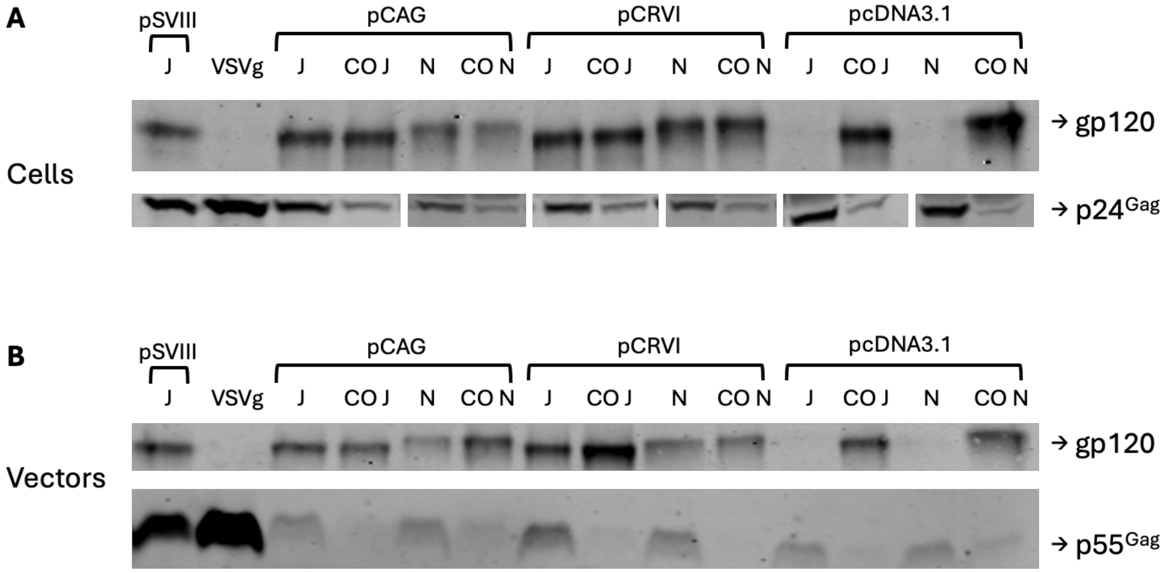


**Figure S2: HIV-1 Env expression and incorporation is similar between pSVIII and pCAG expression plasmid.**
HEK 293T cells were transfected with JR-FL Env being expressed by different plasmids. **(A)** Producer cells and **(B)** vectors were harvested and assessed by immunoblotting for HIV-1 Env and Gag.

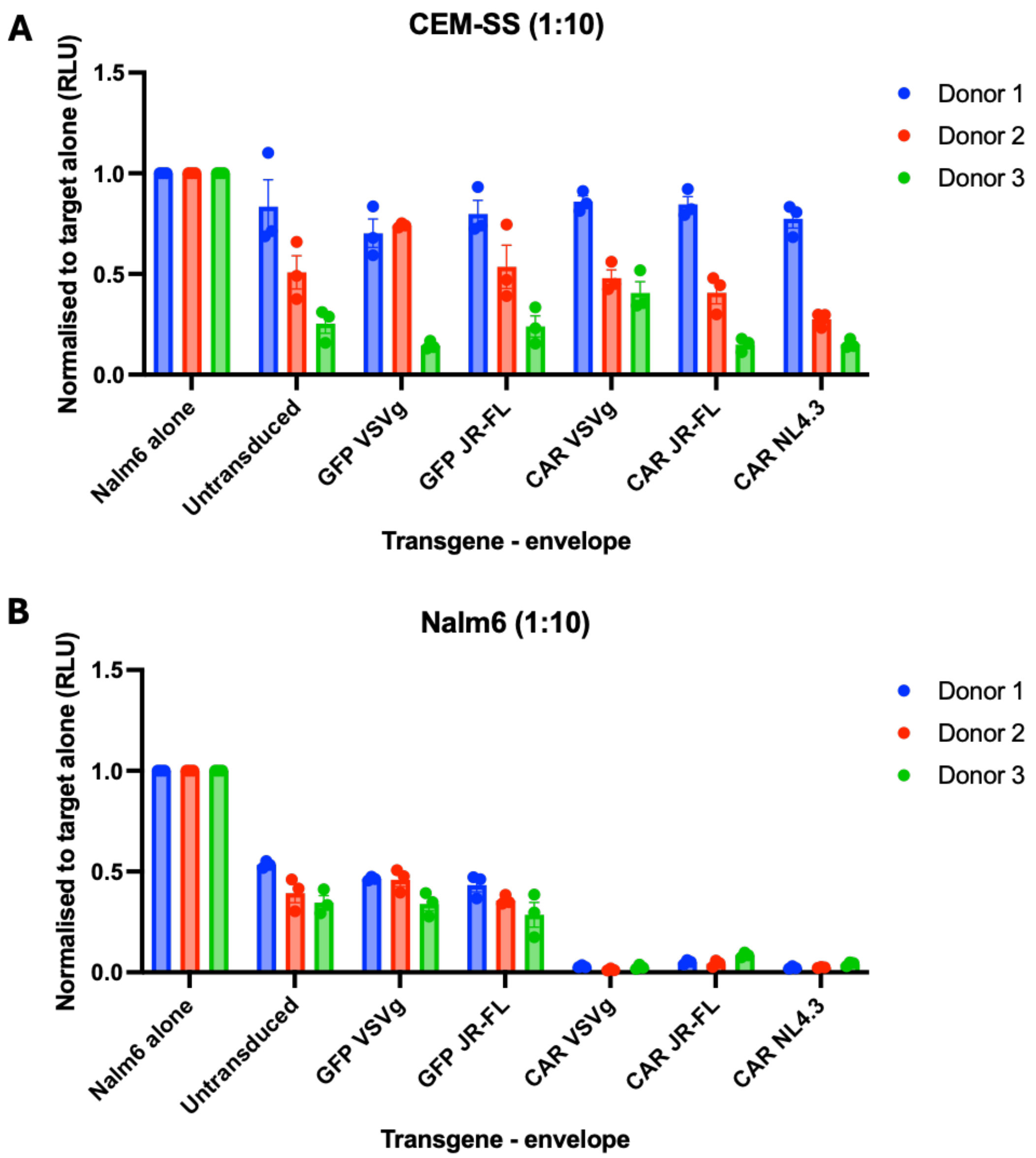


**Figure S3: CAR T cells kill target cells effectively at 1:10 effector to target ratio**
Activated CD4$^{+}$ T cells transduced with vectors with VSV-G and HIV-1 Envs expressing a chimeric antigen receptor (CAR) that targets CD19 or GFP were co-cultured for 48 hours at an effector to target ratio of 1:10 with **(A)** CEM-SS, **(B)** Nalm6. Cytotoxicity was assessed by measuring luciferase activity in surviving cells. Graphs show mean ± SEM of luciferase activity normalised to target cells alone.

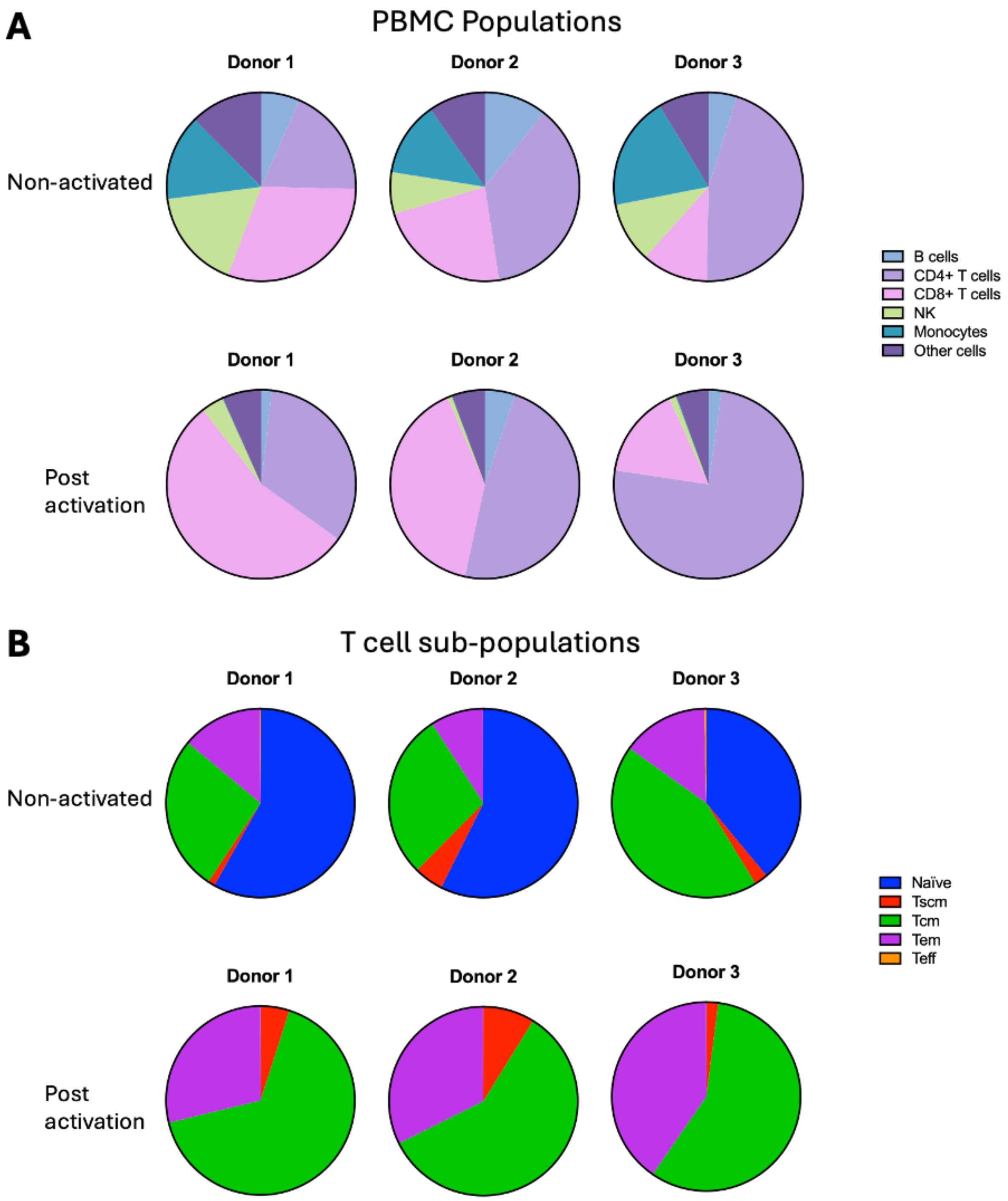


**Figure S4: PBMC activation changes cell population frequencies.**
Pie chart quantification of specific cell populations in PBMCs before (top) and after (bottom) activation with CD3/CD28 beads and IL-2 for 24 hours. Cells were stained with a live/dead dye and cell surface markers and analysed using flow cytometry. Pie charts display the ratios of cell populations detected in **(A)** PBMC populations and **(B)** $CD4^{+}$ T cell sub-populations for each donor.
